# The effects of habitat structure and lighting on object and background appearance

**DOI:** 10.1101/2025.02.19.639053

**Authors:** George R.A. Hancock, Innes C. Cuthill, Jolyon Troscianko

## Abstract

The color, physical structure, and lighting of environments characterize their appearance to animals and, as a result, play an important role in the evolution of animal coloration. Yet despite this long-standing association, how alterations in lighting from atmospheric conditions and three-dimensional structure interact to alter the appearance of scenes and animals remain largely unquantified. This gap is largely due to the lack of controlled measurement of visual properties under different lighting conditions and the formerly limited availability of 3-dimensional (3D) scanning technology. To investigate these interactions, we quantified the appearance of natural backgrounds and a standardized target, with color-calibrated photographs and 3D scans of 672 natural scenes taken under diffuse and direct lighting conditions. We find several instances where lighting and the immediate 3D environment systematically altered the contrast and pattern orientation of scenes and of a target object. Shadows formed under direct lighting drastically increased luminance and short-to-long wave (blue-yellow) contrast across spatial scales, especially at larger spatial scales for habitats with greater 3D variation. Conversely, medium-to-long wave (green-red) information was highly stable to changes in lighting. Direct lighting also increased the directionality of scene appearance and the target, but less so when 3D variation was higher. By quantifying how contrast, size and orientations of patterning within a scene change with habitat structure and lighting conditions, we provide a framework for understanding why different animal color patterns are favored for crypsis, or conspicuousness in different habitats.

## Introduction

The evolution of animal color patterns is strongly influenced by the visual features of their surrounding environment and both the visual systems and positioning of their observers (Endler, 1978; Endler, 1990; Endler, 1991; Endler and Thery, 1996). The appearance of the background of any visual scene an animal is observed in is the product of the material properties of its surfaces and the interactions between its shape (surface geometry) and illumination, typically the sun but also reflections from the sky and ambient objects (Endler, 1990; Nascimento et al., 2016; Párraga et al., 1998; Penacchio et al., 2015). Correlations between an environment’s structure and appearance allow us to readily identify and distinguish habitats, be they deserts or jungles. However, natural scenes are not stable through time, even when the physical features do not change; the intensity, orientation, and directionality of the light source within scenes change with the sun’s position (time of year and day) and when light is scattered by atmospheric particles or cloud cover (Nascimento et al., 2016; Penacchio et al., 2015). Lighting changes interact with the geometry of the whole environment, from the immediate surround of the animal to more distal structures. These interactions should alter the contrast and orientation of shadows at different spatial scales, both across the scene and within individual objects of interest (“targets”), such as animals, fruits, or other resources (Cuthill et al., 2019; Dee and Santos, 2011; Rowe et al., 2024; Ruxton et al., 2004). Alterations in the contrast, size and orientations of patterning within a scene from shadows are important for both quantifying the conspicuousness of signals and camouflaged animals, and for evaluating the visual complexity of a scene (Dimitrova and Merilaita, 2010; Xiao and Cuthill, 2016). For instance, an animal with dark spotted patterns may only match the size and contrast of patterns in the background when under direct lighting within the geometry of its natural habitat, but not when under diffuse illumination.

Most research on geometry and lighting associations between animal and background appearance has focused on broad differences between closed environments, such as tropical and temperate forests, and open environments, such as deserts, grasslands, and mountains, which are characterized by the presence or absence of high tree cover (Allen et al., 2011; Endler, 1993; Vinther et al., 2016). This classification has been used to investigate various aspects of animal ecology, including sensory, navigational, and thermal adaptations (Cheng et al., 2018; Cooney et al., 2022; Napier, 1967; Stuart–Fox and Ord, 2004). Increased geometric complexity of closed habitats has been associated with an increased visual complexity, which increases the difficulty of detecting individual objects and animals compared to open habitats (Anstey, 1963; Dimitrova and Merilaita, 2010; Rowe et al., 2021; Xiao and Cuthill, 2016). Comparisons of animal coloration between habitats have shown that animals in closed environments are more colorful and contrasting in their maculation (surface markings), either to signal or to camouflage (Cooney et al., 2022; Dalrymple et al., 2015; Ortolani, 1999). Furthermore, differences in pattern shape and lighting between these environments are thought to explain differences in the shapes of maculation in camouflaged animals (Allen et al., 2011; Caro, 2005; Somveille et al., 2016). That closed habitats are often associated with more contrasting maculation has been proposed to be due to the formation of more complex patterning from shadows, but with less contrasting countershading thought to be due to the diffusing of light by the forest canopy reducing self-shading contrast (Allen et al., 2011; Allen et al., 2012; Penacchio et al., 2015). Yet how the geometry of these environments influences the appearance of the scene lacks direct quantification, instead relying on subjective classification of habitats on a spatial scale relevant to human beings. Furthermore, how the immediate 3D environment, i.e. the area immediately surrounding the animal, interacts with the visual features of the scene and the animal has yet to be assessed.

Classification of habitats as either open or closed has mostly been from an anthropocentric viewpoint and so using this classification will vary widely in its effects depending on the size, orientation and position of the observer and the target object/animals. For instance, a grassland may be an open habitat to an eagle or human with their elevated viewing angles, but to a weasel or jumping spider it contains multiple obstructions, i.e., it is closed (Hancock et al., 2023). It is the scale of 3D features relative to the observer and target that determines whether a visual scene is open or closed. Similarly to obstruction of vision, obstruction of light can occur from any surrounding surface and in turn affect the scene by producing shadows and by altering the color and intensity of the reflected light due to its material properties. Shadows large enough to be cast onto a target object, referred to here as received shadows, should change the spatial scale, orientation and directionality of its internal luminance contrast (Figure 1). Animal and background patterns come in various shapes and directions. Isotropic patterns have no clear directionality in their visual features, while anisotropic patterns are typified by strong stripes in one direction, usually parallel or perpendicular to the body. Matching the orientation features of the local background has been shown to influence the detection of camouflaged targets (Barnett et al., 2017; Penacchio et al., 2018; Pike, 2018; Troscianko et al., 2017). Some animals with anisotropic patterns orientate their body patterns in the direction of background features to the benefit of camouflage (Hughes et al., 2015; Kang et al., 2012; Ulmer et al., 2013). Depending on a target’s shape, facing direct sunlight produces highly contrasting and directional shadows in the orientation of the light. Bilateral targets that orientate towards the sun are more difficult to detect by birds, potentially due to reducing the size of their cast shadow (Mavrovouna et al., 2021).

**Figure 1.**
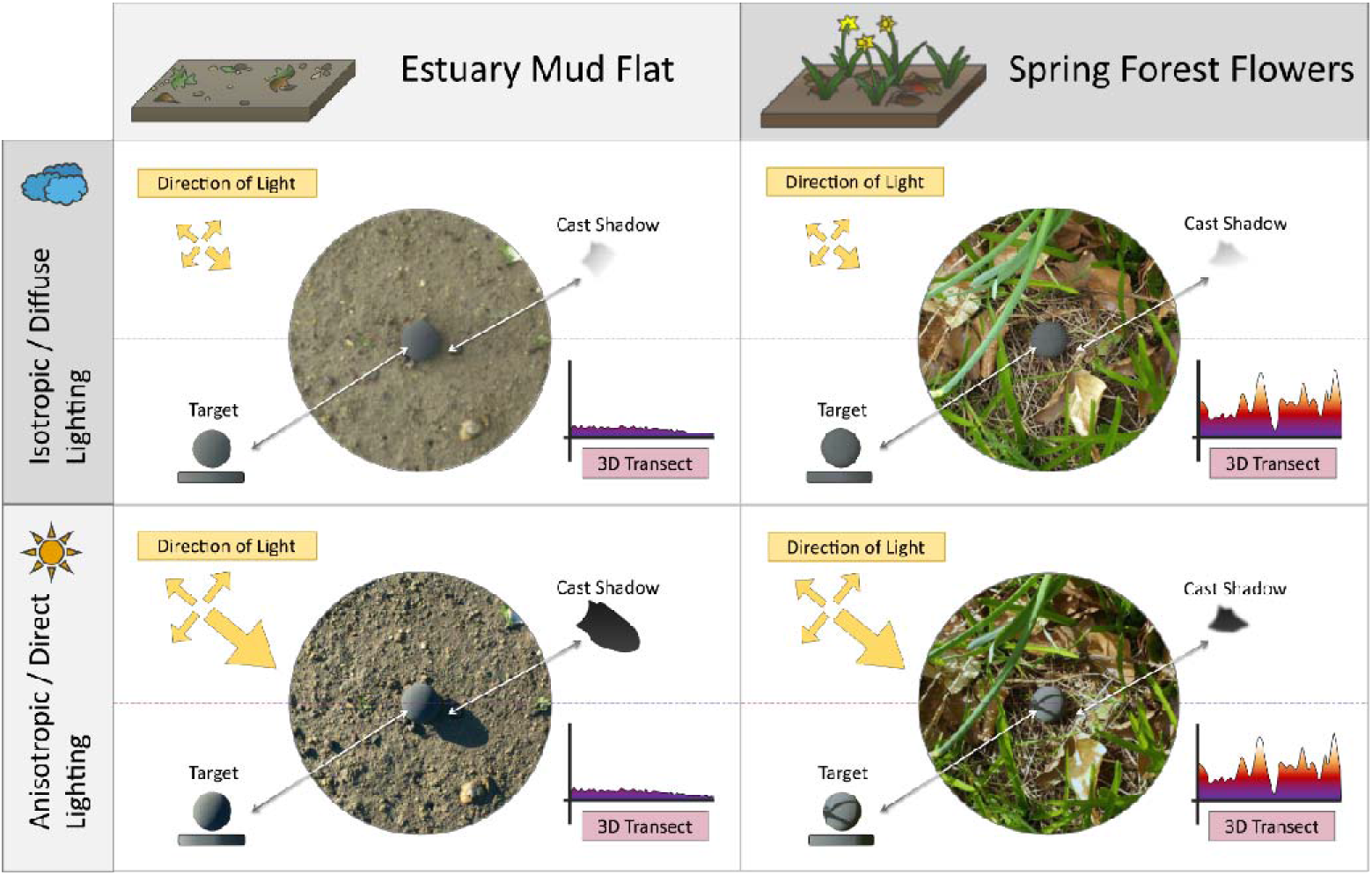
Schematic of interactions between local geometry (physical structure, including depth variation) and weather on the appearance of a target. Plots show depth variation in a 2D cross-section of the 3D scan and so show structure irrespective of lighting. The dashed purple line shows the center line for the 3D transect which extends beyond the image. The top-left shows a target under cloudy conditions with diffuse lighting within a mudflat where the shape and cast shadows are barely visible. The bottom-left shows the same target under sunny conditions with direct lighting, where cast and shape shadows are highly contrasting. The right shows the same target under the two lighting conditions but in a spring forest flower habitat. At the target’s scale, the surrounding geometry of the forest vegetation softens the light, flattening the self-shadow gradient and imposing received shadows perpendicularly to the edge of the self-shadow.

Weather can also affect the scene independently from geometry. On clear sunny days, the high directionality of light produces conspicuous shadows both internal (self) and external (cast) to objects orientated away from the light and specular highlights on surface edges facing the light (Figure 1) (Dee and Santos, 2011; Franklin and Ospina-Rozo, 2021; Kjernsmo et al., 2020). Shadows and highlights increase luminance contrast and, when there is little cloud cover, opponent color contrast due to the blueness of the sky created by Rayleigh scattering (Narasimhan and Nayar, 2002; Steverding and Troscianko, 2004). The increase in longwave (lw) relative to shortwave (sw) light in the direction of the sun makes highlighted regions appear yellower and shadows appear bluer for any animal whose color vision includes lw-sw opponency (blue-yellow contrast in human vision) which encompasses most animals with chromatic sensitivity (Hunt et al., 2009; Kelber and Osorio, 2010; Newport et al., 2017). These effects change under diffuse lighting on overcast days. Here lighting is less intense and more even across directions, reducing contrast and the clarity of shadows (Figure 1). To function effectively, animal coloration often needs to be adapted to changes in illumination from the weather (Penacchio et al., 2018). The common occurrence of red as a signaling color in animals and plants is possibly because of the increased stability of mw-lw contrast compared with sw-lw contrast under changing illumination (Arenas et al., 2014; Lovell et al., 2005). Closed environments in particular are considered more variable in their illumination due to the generation of dynamic lighting from swaying branches and the reduced intensity of light within shadowed regions (Cuthill et al., 2019; Endler, 1993), which itself affects detectability (Matchette et al., 2018; Matchette et al., 2019).

Despite the potential impacts of the directionality of light on how geometry influences the appearance of the scene, thus far its effects have yet to be measured under controlled conditions. Nor has the influence of received shadows on the appearance of target objects. To bridge this gap, we investigated how local geometry and the directionality of lighting influenced the appearance of habitats and target objects. To do so, we artificially manipulated the diffuseness of light within habitats and quantified the visual appearance and 3D structure using color-calibrated photography and hand-held 3D scanners respectively. These methods allowed us to empirically measure how increased variability of depth, “i.e. 3D complexity”, influenced the luminance contrast, chromatic contrast, and the directionality of patterns within the scene at different spatial scales as well as for a target object as a representative model for an animal. Our predictions being that: (1) direct lighting would increase luminance contrast, lw-sw contrast, and the directionality patterns within the scene as well as the internal luminance contrast and directionality of our target object. (2) 3D complexity should increase luminance and chromatic contrast within the scene. (3) Direct lighting should interact with 3D complexity, with more 3D complex scenes producing greater luminance and lw-sw contrasts under direct lighting then less 3D complex, “flatter” scenes, in addition to changing the orientation and scale of contrast on our target due to received shadows from surrounding 3D objects and surfaces.

## Materials and Methods

### (a) Photography

Photographs of 28 temperate habitats were taken within the UK (Figure 2). Habitats were selected to provide a range of geometric and chromatic levels of complexity from a list of targeted landscapes (anthropogenic, grassland, estuaries, forest and heathland). For the anthropogenic habitats, we avoided using completely unnatural materials or structures, such as stone paving and asphalt, which animals rarely use for refuge or foraging. For each habitat, its name, location, time of year, time of day, solar angle, solar azimuth and whether it was classified as a closed (all forest habitats) or open environment were recorded (see supplementary material).

**Figure 2.**
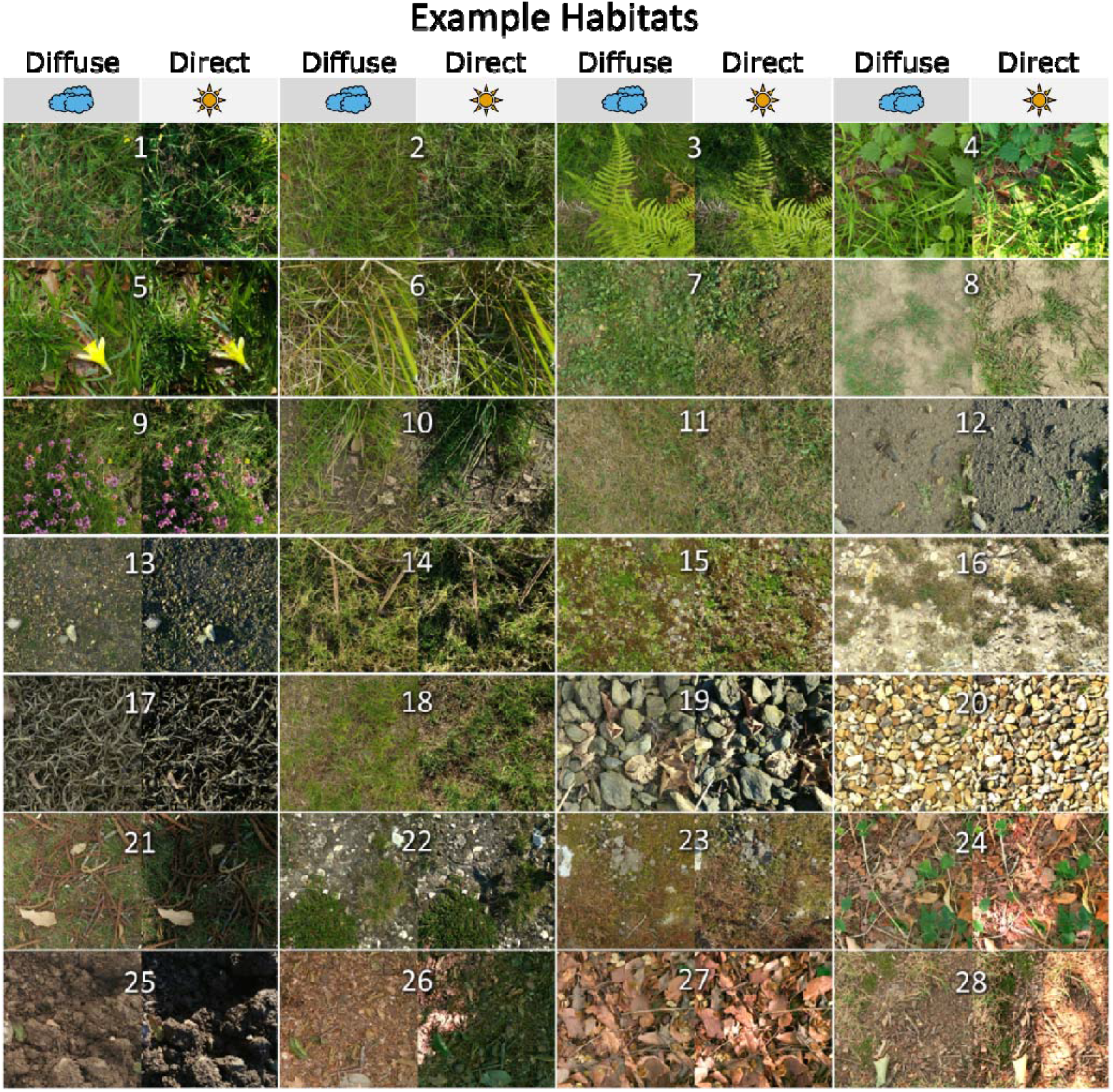
Matrix of samples from background images for all 28 habitats in order of CIE L*a*b* luminance (L*) (left-right) and green-red (a*) (top to bottom). Each background image has been paired with its corresponding photograph under diffuse (left) and direct lighting (right). Closed habitats are denoted using a *. *[1] field wildflower, [2] field topped, [3] ferns, [4] forest nettles*, [5] forest spring flowers*, [6] bog grass, [7] scrub grass, [8] field eroded, [9] heath heather, [10] pond bank, [11], lawn cut, [12] mud flat, [13] estuary gravel, [14] estuary plants, [15] gravel succession, [16] heath eroded, [17] estuary seaweed, [18] lawn uncut, [19] estuary pebbles, [20] gravel, [21] forest succession*, [22] heath chalk, [23] heath succession, [24] forest young bramble*, [25] field plowed, [26] forest trail*, [27] forest leaflitter*, [28] forest edge\**.

For each habitat, 24 scenes (individual locations within the habitat) were photographed in plain view at an angle perpendicular to the ground (birds eye view) under direct and diffuse lighting conditions. This gave a total of 672 scenes across all 28 habitats. Photographs were taken at multiple exposures from a height of 1m using an ASUS Zenfone (ZenFone AR ZS571KL, AsusTek Computer Inc., Taipei, Taiwan). We selected this camera due to its relatively flat field of view, high photo resolution and 3D scanner enabled for Google’s depth-sensor technology ‘Tango’. This scanner works by emitting an infrared grid onto the environment and captures deformation using time-of-flight to produce point clouds.

Photos were taken as non-HDR lossless JPEGs, with a pixel resolution of 5488 × 4096 px and 3D scans were taken using the Matterport Scenes application. To provide direct lighting, all photos were taken on clear days, with less than 10% cloud cover, no obstruction of the sun by cloud and always over two hours from sunrise and sunset. For diffuse lighting, the photographs were repeated with a 1.5m^3^ photography tent placed over the scene (NEEWER, Shenzhen, China). A small rectangular slit was cut into one of the side panels, allowing access to the camera’s horizontal tripod arm (Velbon Ultrek UT-63D Tripod, Hakuba Photo Industry Co., Ltd, Tokyo 130-0014, Japan). The tripod was angled such that neither the shadow of the tripod nor the photographer was visible within the scene.

### (b) Image and Scan Processing

Photographs were calibrated using a hemispheroid “button” grey standard (3D printed with 30mm diameter, spraypainted with Black 3.0 paint mixed with Barium Sulphate to give a diffuse 8% reflectance) (Murray and Adams, 2019). This button acted as a luminance standard and the target object for lighting analyses. The buttons were placed at the center of each scene using a crosshair. A non-overexposed image was selected and linearized for each scene using the ImageJ MICA toolbox (Schneider et al., 2012; Van Den Berg et al., 2020). The grey-target and any occluding vegetation were then labeled as Regions of Interest (ROIs). Post linearization, any vegetation occluding the buttons was manually removed using the clone tool in the open-source image manipulation software GIMP 2.10.36 (GIMP, https://www.gimp.org/; see supplementary material). 3D scans were standardized in MeshLab v.2022.02 and exported as .ply files containing only the X,Y, and Z coordinates. ply files were loaded in ImageJ with a custom script to render an X,Y, Z image where 1px equated to 1mm. Blank z values were replaced locally using nearest-neighbor filling, and the grey-targets were labeled using ROIs (Hancock et al., 2023).

### (c) Measurements

For each scene, the color images and height maps were rescaled using the minimum number of pixels per mm. The color images were converted to the CIE L*a*b* color space using ImageJ’s built-in Lab converter, which divides the image into a luminance channel (L*) and two opponent channels, green-red (a*) and blue-yellow (b*) (CIE, 1976; Renoult et al., 2017). Although CIE L*a*b* is human-specific, it does the important job, common to all general-purpose color vision systems, of decorrelating the photoreceptor outputs to create approximately orthogonal sources of information (Buchsbaum and Gottschalk, 1983; Ruderman et al., 1998). Certainly the large variation in photoreceptor complement and retinal organization across species (Baden et al., 2020) will mean that some of our conclusions will not generalize in all cases. The most obvious limitation is that our camera was unable to photograph in UV and so color opponent channels that efficiently code the chromatic information available to mammals will not be appropriate for insects (Peitsch et al., 1992) lack some of the information available to vertebrates such as birds (Cuthill et al., 2000). However, we are confident that our analyses allow broad conclusions to be drawn about effect on perceptions of variation in overall light intensity, as captured by luminance, and the relative proportions of short-, medium- and longwave light (see e.g. Endler 1990). We also note that the influence of Rayleigh scattering on uv-to-lw opponency is likely to be similar, probably exaggerated, compared to sw-to-lw (Tedore and Nilsson, 2019). In terms of spatial information, many earlier studies on color and luminance patterns within natural scenes have used Fourier transformation of images at different spatial frequencies; here, we opted to use Gabor filters at different spatial scales relative to the target’s diameter, allowing for comparison of angular differences in pattern across multiple spatial scales (See Supplementary Figure A4) (Párraga et al., 1998; Talas et al., 2017). Gabor filters work by using orientation-selective bandpass filters that mirror the spatial receptive fields of early visual processing (Field, 1987, 19).

For each channel (L*, a*, b*) the mean was measured, as well as the contrast (Standard Deviation, StdDev) for the scene filtered for 8 different spatial scales [0.878mm, 1.758mm, 3.516mm, 7.031mm, 14.062mm, 28.125mm, 56.25mm, 112.5mm] and 4 different gabor angles [0°,45°,90°,135°], where 0° was parallel with the y axis and 90° with the x axis (Barnett et al., 2021; Pike, 2018). Scales larger than the target diameter (30mm) were excluded from the target analyses. The contrast at a given spatial scale was measured as the mean contrast of all angles. Directionality was measured for each scale by dividing the angle with the highest contrast by the mean contrast of the other angles. The further the value was from 1, the lower the isotropy and the greater the level of directionality. For scenes under direct lighting, pattern contrast parallel [0°] to and perpendicular [90°] to the direction of light of the target was also measured as an estimation for received and self-shading, respectively by re-orientating the target to match the direction of the encroaching light. The local 3D complexity of the environment was measured using the local depth variation (StdDev of Z values) of the 3D scans. Sample calibrated scene images and scans are included in our supplementary material along with the custom image analysis scripts for ImageJ.

### Statistical Analyses

Statistical analyses were performed using R, version 4.3.0 (R Core Team, 2023). The metrics that were expected to be influenced by local depth variation and lighting conditions (luminance contrast, luminance directionality, blue-yellow contrast, red-green contrast, and pattern orientations for the target) were tested with the lme4 package using linear mixed models (Bates et al., 2014). Each background and target metric was given as the dependent variable, with depth variation, the lighting type and the spatial scale as the predictor variables. The unique ID for each of the 672 scenes was used as a random effect to control for the correlation in measures at a given location under the two lighting conditions. Given that shadows were likely to influence contrasts and directionality differently across scales, but not necessarily linearly, we used Akaike information criterion (AIC) analysis to determine whether a linear model or a polynomial model, either quadratic or cubic, for spatial scale was most appropriate. For all analyses a cubic model was the best fit (lowest AIC value), an example formula is given below:

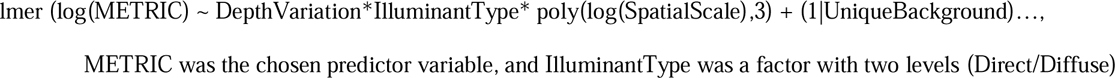

Tested metrics were as follows for the backgrounds: log(luminance contrast), log(luminance directionality), log(blue-yellow contrast) and log(red-green contrast). For the grey-targets: log(luminance contrast), log(luminance directionality), were measured for all targets, as well as the proportion of contrast oriented parallel and perpendicular to the angle of light (angle contrast/sum contrast), but only for the directly illuminated targets. Log transformations were used to ensure normality of our data (Stevens et al., 2014). For the orientation analysis, the linear models were modified to include habitat categorization as a closed environment, i.e., the woodland habitats, instead of illuminant type, or open habitat, as canopy cover is known to influence self-shading and measurements were only taken under the direct illumination treatment. To meet the assumptions of our models, the residuals were checked for normality and homogeneity of variance.

## Results

To simplify the breadth of interactions, tabulated model outputs are provided for the background (Tables 1 and 2) and grey-target statistics (Table 3).

**Table 1.**
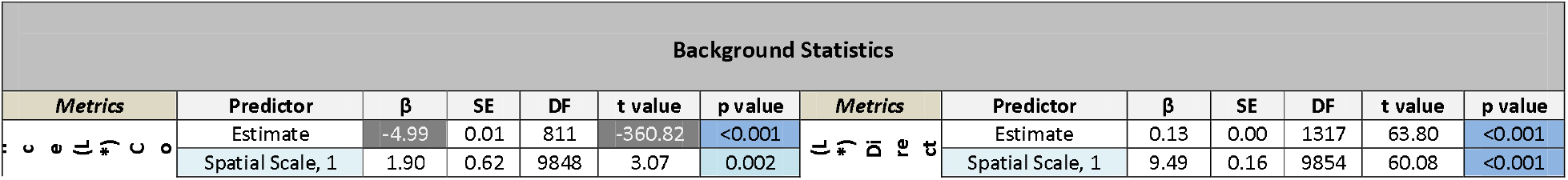
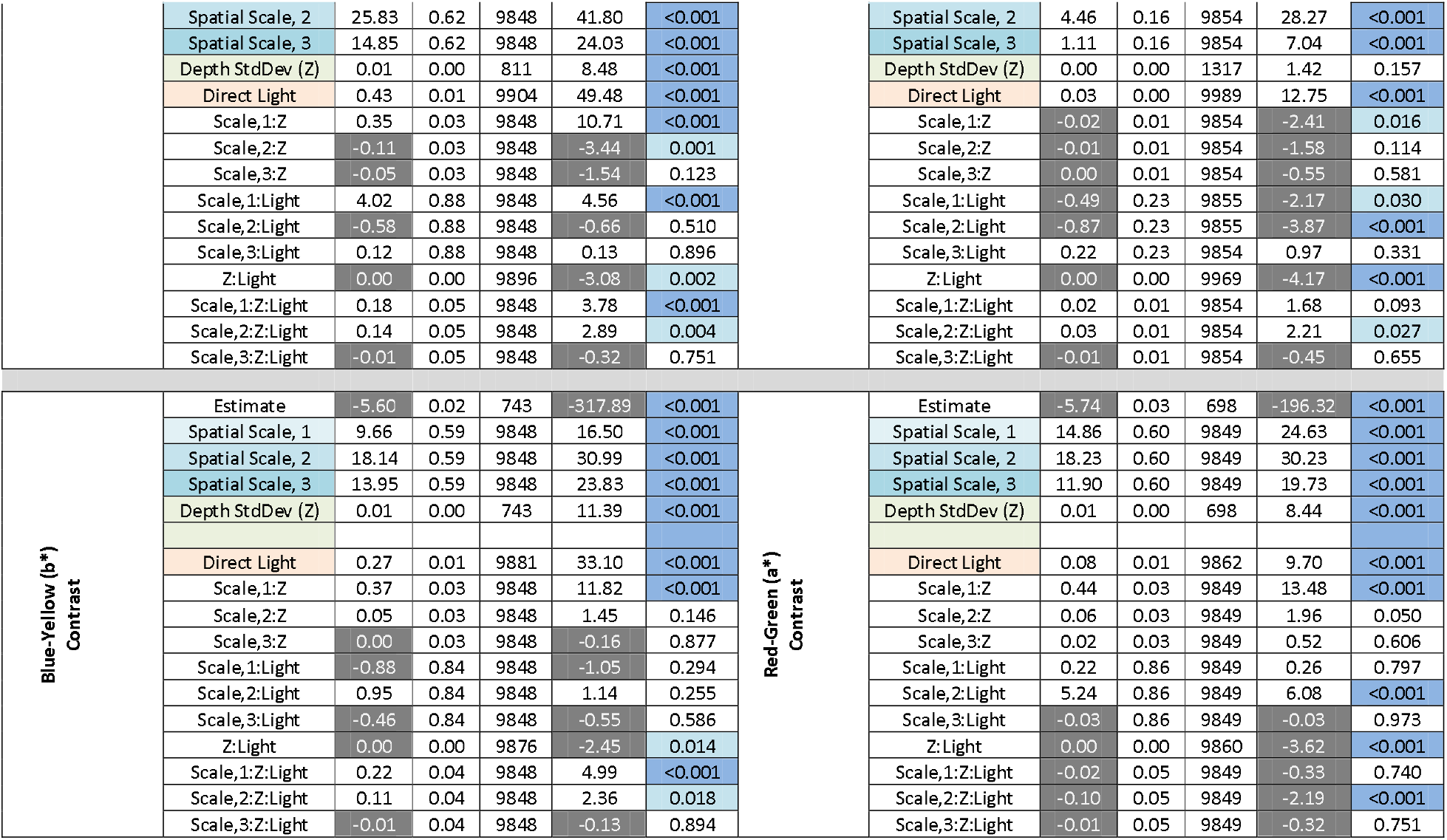
LMM results for interactions between spatial scale, background depth variation (Z StdDev), the directionality of illumination and background statistics with the estimate (β), standard error (SE), degrees of freedom (DF), t-value and p-value for each interaction. Instances where the effects were significant (p<0.05),) are marked in blue. Positive estimates and t-values are shown against a white background, while negative values are shown against dark grey. For each analysis, the effect of spatial scale is split using a 3-part polynomial where the number indicates the interaction level (1=linear term, 2=quadratic term, and 3= cubic term).

| Background Statistics |  |  |  |  |  |  |  |  |  |  |  |  |  |
| *Metrics* | Predictor | β | SE | DF | t value | p value | *Metrics* | Predictor | β | SE | DF | t value | p value |
|---|---|---|---|---|---|---|---|---|---|---|---|---|---|
| [illegible] | Estimate | -4.99 | 0.01 | 811 | -360.82 | <0.001 | [illegible] | Estimate | 0.13 | 0.00 | 1317 | 63.80 | <0.001 |
|  | Spatial Scale, 1 | 1.90 | 0.62 | 9848 | 3.07 | 0.002 |  | Spatial Scale, 1 | 9.49 | 0.16 | 9854 | 60.08 | <0.001 |
|  | Spatial Scale, 2 | 25.83 | 0.62 | 9848 | 41.80 | <0.001 |  | Spatial Scale, 2 | 4.46 | 0.16 | 9854 | 28.27 | <0.001 |
|  | Spatial Scale, 3 | 14.85 | 0.62 | 9848 | 24.03 | <0.001 |  | Spatial Scale, 3 | 1.11 | 0.16 | 9854 | 7.04 | <0.001 |
|  | Depth StdDev (Z) | 0.01 | 0.00 | 811 | 8.48 | <0.001 |  | Depth StdDev (Z) | 0.00 | 0.00 | 1317 | 1.42 | 0.157 |
|  | Direct Light | 0.43 | 0.01 | 9904 | 49.48 | <0.001 |  | Direct Light | 0.03 | 0.00 | 9989 | 12.75 | <0.001 |
|  | Scale,1:Z | 0.35 | 0.03 | 9848 | 10.71 | <0.001 |  | Scale,1:Z | -0.02 | 0.01 | 9854 | -2.41 | 0.016 |
|  | Scale,2:Z | -0.11 | 0.03 | 9848 | -3.44 | 0.001 |  | Scale,2:Z | -0.01 | 0.01 | 9854 | -1.58 | 0.114 |
|  | Scale,3:Z | -0.05 | 0.03 | 9848 | -1.54 | 0.123 |  | Scale,3:Z | 0.00 | 0.01 | 9854 | -0.55 | 0.581 |
|  | Scale,1:Light | 4.02 | 0.88 | 9848 | 4.56 | <0.001 |  | Scale,1:Light | -0.49 | 0.23 | 9855 | -2.17 | 0.030 |
|  | Scale,2:Light | -0.58 | 0.88 | 9848 | -0.66 | 0.510 |  | Scale,2:Light | -0.87 | 0.23 | 9855 | -3.87 | <0.001 |
|  | Scale,3:Light | 0.12 | 0.88 | 9848 | 0.13 | 0.896 |  | Scale,3:Light | 0.22 | 0.23 | 9854 | 0.97 | 0.331 |
|  | Z:Light | 0.00 | 0.00 | 9896 | -3.08 | 0.002 |  | Z:Light | 0.00 | 0.00 | 9969 | -4.17 | <0.001 |
|  | Scale,1:Z:Light | 0.18 | 0.05 | 9848 | 3.78 | <0.001 |  | Scale,1:Z:Light | 0.02 | 0.01 | 9854 | 1.68 | 0.093 |
|  | Scale,2:Z:Light | 0.14 | 0.05 | 9848 | 2.89 | 0.004 |  | Scale,2:Z:Light | 0.03 | 0.01 | 9854 | 2.21 | 0.027 |
|  | Scale,3:Z:Light | -0.01 | 0.05 | 9848 | -0.32 | 0.751 |  | Scale,3:Z:Light | -0.01 | 0.01 | 9854 | -0.45 | 0.655 |
| **Blue-Yellow (b\*)<br>Contrast** | Estimate | -5.60 | 0.02 | 743 | -317.89 | <0.001 | **Red-Green (a\*)<br>Contrast** | Estimate | -5.74 | 0.03 | 698 | -196.32 | <0.001 |
|  | Spatial Scale, 1 | 9.66 | 0.59 | 9848 | 16.50 | <0.001 |  | Spatial Scale, 1 | 14.86 | 0.60 | 9849 | 24.63 | <0.001 |
|  | Spatial Scale, 2 | 18.14 | 0.59 | 9848 | 30.99 | <0.001 |  | Spatial Scale, 2 | 18.23 | 0.60 | 9849 | 30.23 | <0.001 |
|  | Spatial Scale, 3 | 13.95 | 0.59 | 9848 | 23.83 | <0.001 |  | Spatial Scale, 3 | 11.90 | 0.60 | 9849 | 19.73 | <0.001 |
|  | Depth StdDev (Z) | 0.01 | 0.00 | 743 | 11.39 | <0.001 |  | Depth StdDev (Z) | 0.01 | 0.00 | 698 | 8.44 | <0.001 |
|  | Direct Light | 0.27 | 0.01 | 9881 | 33.10 | <0.001 |  | Direct Light | 0.08 | 0.01 | 9862 | 9.70 | <0.001 |
|  | Scale,1:Z | 0.37 | 0.03 | 9848 | 11.82 | <0.001 |  | Scale,1:Z | 0.44 | 0.03 | 9849 | 13.48 | <0.001 |
|  | Scale,2:Z | 0.05 | 0.03 | 9848 | 1.45 | 0.146 |  | Scale,2:Z | 0.06 | 0.03 | 9849 | 1.96 | 0.050 |
|  | Scale,3:Z | 0.00 | 0.03 | 9848 | -0.16 | 0.877 |  | Scale,3:Z | 0.02 | 0.03 | 9849 | 0.52 | 0.606 |
|  | Scale,1:Light | -0.88 | 0.84 | 9848 | -1.05 | 0.294 |  | Scale,1:Light | 0.22 | 0.86 | 9849 | 0.26 | 0.797 |
|  | Scale,2:Light | 0.95 | 0.84 | 9848 | 1.14 | 0.255 |  | Scale,2:Light | 5.24 | 0.86 | 9849 | 6.08 | <0.001 |
|  | Scale,3:Light | -0.46 | 0.84 | 9848 | -0.55 | 0.586 |  | Scale,3:Light | -0.03 | 0.86 | 9849 | -0.03 | 0.973 |
|  | Z:Light | 0.00 | 0.00 | 9876 | -2.45 | 0.014 |  | Z:Light | 0.00 | 0.00 | 9860 | -3.62 | <0.001 |
|  | Scale,1:Z:Light | 0.22 | 0.04 | 9848 | 4.99 | <0.001 |  | Scale,1:Z:Light | -0.02 | 0.05 | 9849 | -0.33 | 0.740 |
|  | Scale,2:Z:Light | 0.11 | 0.04 | 9848 | 2.36 | 0.018 |  | Scale,2:Z:Light | -0.10 | 0.05 | 9849 | -2.19 | <0.001 |
|  | Scale,3:Z:Light | -0.01 | 0.04 | 9848 | -0.13 | 0.894 |  | Scale,3:Z:Light | -0.01 | 0.05 | 9849 | -0.32 | 0.751 |

### Background Statistics

Direct lighting significantly increased luminance contrast, chromatic contrast and directionality compared to diffuse lighting, with the greatest increase in contrast being in the luminance channel followed by blue-yellow (Figure 3, Table 1). Increased depth variation also correlated with luminance and chromatic contrast at larger spatial scales, although it marginally reduced the degree that direct lighting increased contrast at smaller spatial scales but increased the effect at larger scales. Directionality was higher under direct lighting, but only at smaller spatial scales and was greater when local 3D complexity was lower. For the luminance channel, these changes equate to 3D complexity and direct lighting increasing the steepness of the Fourier slope.

**Figure 3.**
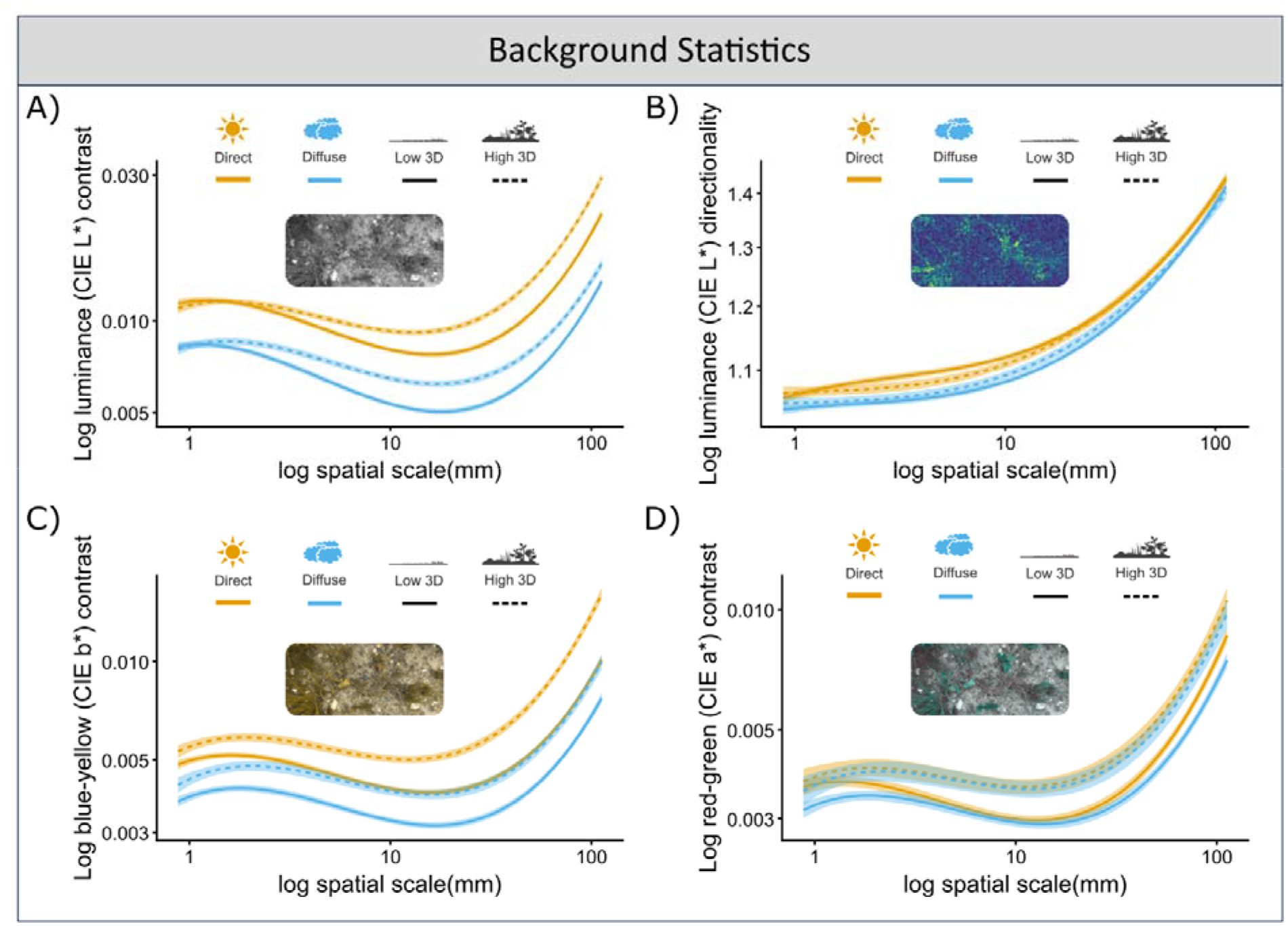
Interactions between spatial scale, background geometry, lighting treatment and background scene statistics. (A) luminance contrast, (B) pattern directionality, (C) blue-yellow contrast and (D) green-red contrast. Channels correspond with CIE L*a*b* color space. Points indicate the value for each scene under the different lighting condition, while the lines depict the Cubic equation fitted for each line, with diffuse shown in blue and direct lighting shown in yellow. The 3D variation is split by line type for depth variation (Z StdDev), with low variation as a smooth line and high variation as a dashed line.

### Target Statistics

As expected, given the hemispheroid shape of the target, luminance contrast peaked at ∼1/2 the diameter of the target (Figure 4A, Table 3). Like the background, the contrast and directionality of luminance increased under direct lighting. However, unlike the background 3D complexity increased contrast and directionality at smaller spatial scales and decreased directionality at larger scales under direct lighting. Closed (woodland) environments had significantly lower proportions of contrast perpendicular to the light than (self-shading contrast) open environments at larger spatial scales. However, increased local depth variation resulted in increased proportions of contrast parallel to the direction of light irrespective of whether the habitat was open or closed, especially at smaller spatial scales.

**Figure 4.**
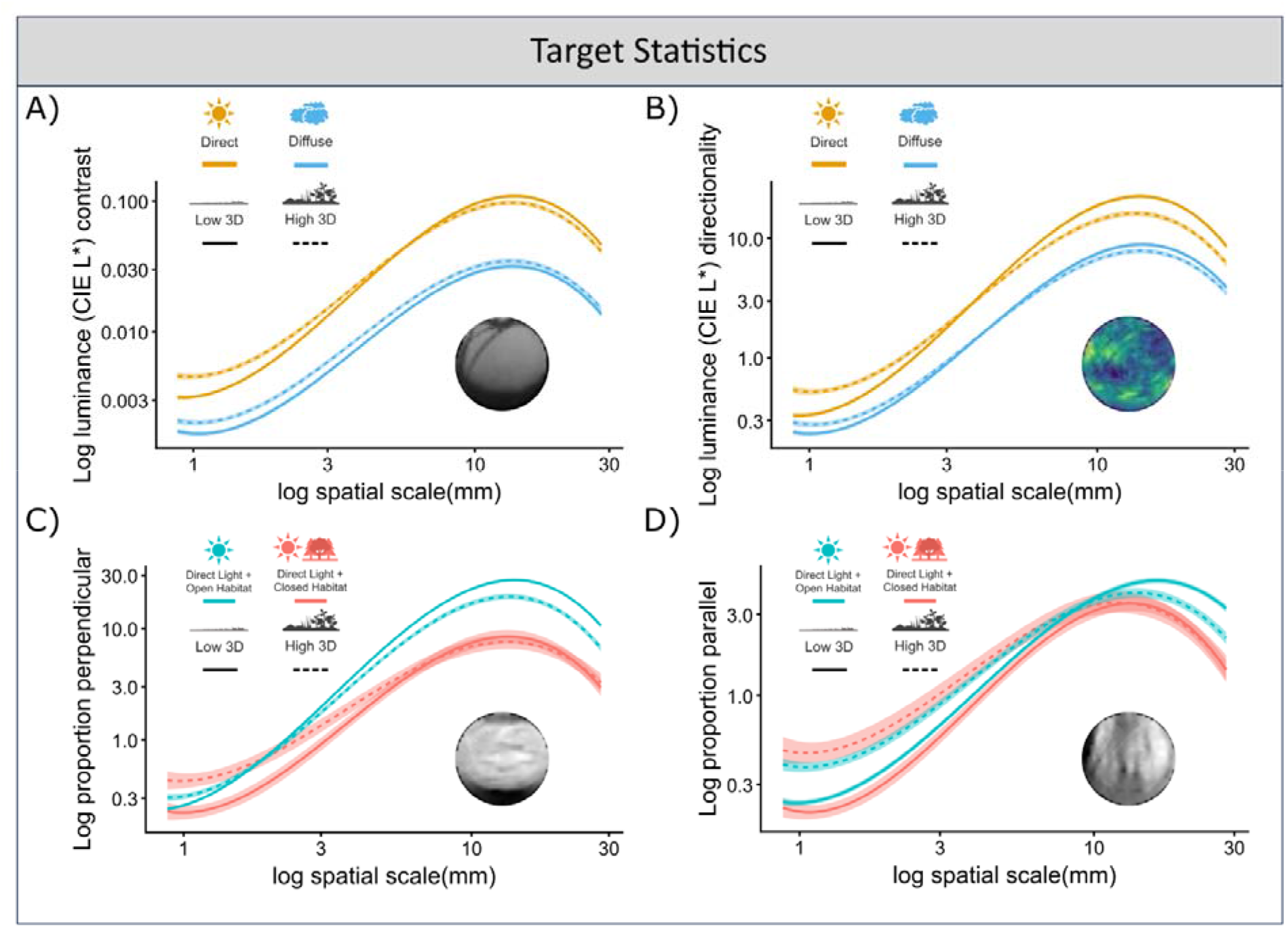
Interactions between spatial scale, background geometry, lighting treatment and target luminance statistics. (A) luminance contrast, (B) pattern directionality, (C) proportion of contrast perpendicular to the direct light source and (D) proportion of contrast parallel to the direct light source. Here luminance is L* within the CIE L*a*b* color space. For the top plots (A and B) lighting is split by the line color, with diffuse shown in blue and direct lighting shown in yellow. For the below plots (C and D) whether the habitat was open (not-woodland) or closed (woodland) is split by line color with open habitats shown in blue and closed habitats in red. Points show the individual values for the target within each scene. For the below plots only photographs under direct lighting were used. The 3D variation is split by line type for depth variation (Z StdDev), with low variation as a smooth line and high variation as a dashed line.

**Table 2.** LMM results for interactions between spatial scale, background depth variation (Z StdDev), the directionality of illumination and target statistics with the estimate (β), standard error (SE), degrees of freedom (DF), t-value and p-value for each interaction. Instances where the effects were significant (p<0.05),) are marked in blue. Positive estimates and t-values are shown against a white background, while negative values are shown against dark grey. Analysis of luminance pattern orientation (perpendicular and parallel) was carried out only for targets under direct lighting. For each analysis, the effect of spatial scale is split using a 3-part polynomial where the number indicates the interaction level (1=linear term, 2=quadratic term, and 3= cubic term).

| Target Statistics |  |  |  |  |  |  |  |  |  |  |  |  |  |
|---|---|---|---|---|---|---|---|---|---|---|---|---|---|
| ***Metrics*** | **Parameter** | **β** | **SE** | **DF** | **t value** | **p value** | ***Metrics*** | **Predictor** | **β** | **SE** | **DF** | **t value** | **p value** |
| **Luminance (L\*)<br>Contrast** | Intercept | -4.89 | 0.02 | 846 | -240.74 | <0.001 | **Luminance (L\*)<br>Directionality** | Intercept | 0.48 | 0.02 | 846 | 20.61 | <0.001 |
|  | Spatial Scale, 1 | 86.75 | 0.87 | 7214 | 99.67 | <0.001 |  | Spatial Scale, 1 | 112.60 | 0.98 | 7216 | 114.49 | <0.001 |
|  | Spatial Scale, 2 | -30.43 | 0.87 | 7214 | -34.96 | <0.001 |  | Spatial Scale, 2 | -32.93 | 0.98 | 7216 | -33.48 | <0.001 |
|  | Spatial Scale, 3 | -34.72 | 0.87 | 7214 | -39.90 | <0.001 |  | Spatial Scale, 3 | -38.23 | 0.98 | 7216 | -38.88 | <0.001 |
|  | Depth StdDev (Z) | 0.01 | 0.00 | 846 | 5.08 | <0.001 |  | Depth StdDev (Z) | 0.00 | 0.00 | 846 | -0.23 | 0.817 |
|  | Direct Light | 1.06 | 0.01 | 7264 | 75.32 | < 2e-16 |  | Direct Light | 0.71 | 0.02 | 7265 | 44.83 | <0.001 |
|  | Scale,1:Z | -0.28 | 0.05 | 7214 | -6.04 | <0.001 |  | Scale,1:Z | -0.54 | 0.05 | 7216 | -10.33 | <0.001 |
|  | Scale,2:Z | -0.03 | 0.05 | 7214 | -0.73 | 0.464 |  | Scale,2:Z | 0.01 | 0.05 | 7216 | 0.12 | 0.902 |
|  | Scale,3:Z | 0.08 | 0.05 | 7214 | 1.75 | 0.081 |  | Scale,3:Z | 0.07 | 0.05 | 7216 | 1.27 | 0.206 |
|  | Scale,1:Light | 21.25 | 1.24 | 7214 | 17.12 | <0.001 |  | Scale,1:Light | 18.23 | 1.40 | 7216 | 13.00 | <0.001 |
|  | Scale,2:Light | -8.64 | 1.24 | 7214 | -6.96 | <0.001 |  | Scale,2:Light | -11.00 | 1.40 | 7216 | -7.84 | <0.001 |
|  | Scale,3:Light | -1.09 | 1.24 | 7214 | -0.88 | 0.380 |  | Scale,3:Light | -3.74 | 1.40 | 7216 | -2.66 | 0.008 |
|  | Z:Light | 0.00 | 0.00 | 7256 | -6.17 | <0.001 |  | Z:Light | 0.00 | 0.00 | 7258 | -2.10 | 0.036 |
|  | Scale,1:Z:Light | -0.44 | 0.07 | 7214 | -6.63 | <0.001 |  | Scale,1:Z:Light | -0.56 | 0.08 | 7216 | -7.50 | <0.001 |
|  | Scale,2:Z:Light | 0.20 | 0.07 | 7214 | 3.05 | 0.002 |  | Scale,2:Z:Light | 0.21 | 0.08 | 7216 | 2.80 | 0.005 |
|  | Scale,3:Z:Light | 0.04 | 0.07 | 7214 | 0.59 | 0.557 |  | Scale,3:Z:Light | 0.08 | 0.08 | 7216 | 1.00 | 0.315 |
| **Proportion<br>Perpendicular** | Intercept | 0.41 | 0.05 | 641 | 7.89 | <0.001 | **Proportion<br>Parallel** | Intercept | -0.15 | 0.05 | 641 | -3.03 | 0.003 |
|  | Spatial Scale, 1 | 76.09 | 1.27 | 3213 | 60.10 | <0.001 |  | Spatial Scale, 1 | 57.11 | 1.39 | 3213 | 41.02 | <0.001 |
|  | Spatial Scale, 2 | -27.93 | 1.27 | 3213 | -22.06 | <0.001 |  | Spatial Scale, 2 | -21.44 | 1.39 | 3213 | -15.40 | <0.001 |
|  | Spatial Scale, 3 | -28.99 | 1.27 | 3213 | -22.90 | <0.001 |  | Spatial Scale, 3 | -25.02 | 1.39 | 3213 | -17.97 | <0.001 |
|  | Depth StdDev (Z) | 0.01 | 0.00 | 641 | 2.81 | 0.005 |  | Depth StdDev (Z) | 0.01 | 0.00 | 641 | 4.76 | <0.001 |
|  | Open Habitat | 0.86 | 0.06 | 641 | 14.45 | < 2e-16 |  | Open Habitat | 0.26 | 0.06 | 641 | 4.65 | <0.001 |
|  | Scale,1:Z | -0.58 | 0.05 | 3213 | -10.67 | < 2e-16 |  | Scale,1:Z | -0.56 | 0.06 | 3213 | -9.40 | <0.001 |
|  | Scale,2:Z | 0.08 | 0.05 | 3213 | 1.48 | 0.140 |  | Scale,2:Z | 0.01 | 0.06 | 3213 | 0.15 | 0.885 |
|  | Scale,3:Z | 0.19 | 0.05 | 3213 | 3.60 | <0.001 |  | Scale,3:Z | 0.16 | 0.06 | 3213 | 2.64 | 0.008 |
|  | Scale,1:Open | 25.23 | 1.45 | 3213 | 17.35 | < 2e-16 |  | Scale,1:Open | 14.82 | 1.60 | 3213 | 9.27 | <0.001 |
|  | Scale,2:Open | -7.75 | 1.45 | 3213 | -5.33 | 0.000 |  | Scale,2:Open | 9.95 | 1.60 | 3213 | 6.22 | <0.001 |
|  | Scale,3:Open | 2.19 | 1.45 | 3213 | 1.50 | 0.133 |  | Scale,3:Open | 6.67 | 1.60 | 3213 | 4.17 | <0.001 |
|  | Z:Open | -0.01 | 0.00 | 641 | -4.61 | 0.000 |  | Z:Open | 0.00 | 0.00 | 641 | -1.58 | 0.114 |
|  | Scale,1:Z:Open | -0.08 | 0.07 | 3213 | -1.22 | 0.223 |  | Scale,1:Z:Open | -0.27 | 0.08 | 3213 | -3.62 | <0.001 |
|  | Scale,2:Z:Open | -0.02 | 0.07 | 3213 | -0.23 | 0.817 |  | Scale,2:Z:Open | -0.20 | 0.08 | 3213 | -2.70 | 0.007 |
|  | Scale,3:Z:Open | -0.22 | 0.07 | 3213 | -3.19 | 0.001 |  | Scale,3:Z:Open | -0.16 | 0.08 | 3213 | -2.07 | 0.039 |

## Discussion

In this study, we demonstrate that changes in the directionality of lighting, such as those that would occur from atmospheric conditions, systematically altered our image-derived metrics for our grey targets and the backgrounds across multiple habitats. Moreover, the role of lighting varies with habitat geometry, affecting both the stability of the visual scene, under variable illumination, and the scale and orientation of pattern changes. Conversely, the geometry of scenes correlated with the scale and contrast of its luminance and color features irrespective of lighting condition. Given the size of the target and the backgrounds we studied, our results are most relevant to small terrestrial animals.

In our study, direct lighting increased contrast caused by self-shading and cast shadows. Depending on the spatial scale, increased surface variation created distorted and overlapping shadows within our scenes, reducing the increase in contrast under direct light but still resulting in greater luminance contrasts overall (Cavanagh and Leclerc, 1989; Dee and Santos, 2011). As none of our backgrounds were completely flat, direct lighting produced directional shadows changing the pattern shape, not just the contrast of the background, especially within flatter backgrounds where the shadows were unbroken. The increased conspicuousness of shadows cast by objects against flatter backgrounds is likely to be consistent across all scales, e.g., the increased conspicuousness of dappled shadows on a flat forest floor compared to on a fern-covered understory or within the canopy itself, but requires quantification in larger scenes.

As our target object was smooth, under direct lighting surrounding vegetation frequently created directional received shadows parallel to the light source. These shadows affected both the internal contrast and the direction of the target’s internal shading at different spatial scales. Determining the effect of lighting on an animal and its background’s appearance thus requires knowledge of the surrounding geometry at scales both similar in size and larger than the animal. The dark maculation of animals has often been observed to resemble received and surrounding cast shadows within their environments created by direct lighting (Armbruster and Page, 1996; Ortolani, 1999; Pembury Smith and Ruxton, 2020). These patterns, depending on their intensity and orientation, may also mask internal contrast and orientation cues of geometry from self-shading (Egan et al., 2016; Kelley et al., 2022; Webster, 2015). Whether received shadows reduce the conspicuousness of self-shading to observers, however, has yet to be quantified and requires behavioral experiments.

Consistent with experiments on single scenes (Lovell et al., 2005) or simple backgrounds (Arenas et al., 2014), green-red contrast within our scenes was distinguished from luminance and blue-yellow contrast by its stability under direct lighting. Instead, green-red contrast was mostly correlated with increased depth variation, likely due to the presence of vegetation (Potetz and Lee, 2003). Previous work on opponent color channels and the statistics of visual scenes have shown that green-red edges do not correlate highly with luminance or blue-yellow edges and the spatial scale of green-red features is often larger than luminance features (Hansen and Gegenfurtner, 2009). The stability of green-red contrast within visual scenes not only makes it an effective cue for object detection, but its correlation with the geometry of vegetation can make it useful for predicting depth when navigating terrestrial 3D scenes and is used as such for computer vision (Kirk et al., 2009; Troscianko et al., 1991). The correlation between 3D complexity and red-green contrast within our images supports this relationship, and may present an advantage to animals with red-green opponency present in most non-mammalian tetrapods, reef-fish, and various flying insects (Newport et al., 2017; Osorio et al., 1999; Przyrembel et al., 1995).

Numerous studies focusing on the efficacy of animal camouflage and signaling colors have photographed target animals and their backgrounds under standardized, usually diffuse or direct flash photography, lighting conditions, with the animal frequently removed from the local background to increase the ease of photography and measurement (Hulse et al., 2020; Stevens et al., 2007). This is with good reason, as shadows can interfere with the measurement of animal patterns and features, making them difficult to measure and compare accurately if lighting is not controlled (Finlayson et al., 2002). However, our results highlight several circumstances where discrepancies in the light environment (sum of geometry and weather) would change the level of measured match between the target and its background (Akkaynak et al., 2017; Hulse et al., 2020). Caution should especially be taken when assessing the efficacy of camouflage and signals based on differences in the scale, contrast and orientation of luminance and long-to-short wave opponent patterns. Animal patterns that appear to be more contrasting than their background under diffuse lighting may match the pictorial contrast of the background or be concealed by self-shading when under direct lighting or when an animal changes its orientation relative to the light. Conversely, patterns that match the contrast of the background may not match in directionality or orientation (Stuart–Fox and Ord, 2004).

In line with prior assumptions of habitat appearance, more locally 3D complex habitats did lower the effect of light on the directionality of the background (Somveille et al., 2016); however, depth variation also increased directionality under diffuse conditions and increased directionality of patterns internal to the target at small spatial scales. Many animals possess directional patterns such as stripes and bars, thought to mimic directional background features. Past analyses of avian and mammalian maculation patterns have found inconsistencies in the predicted and observed shape and orientations of animal patterns within open and closed habitats (Allen et al., 2011; Somveille et al., 2016). Within our study, the level of directionality and pattern contrast within backgrounds was inconsistent across scales and fluctuated with lighting conditions and local geometry. Even in habitats that are primarily flat, stripes could still be background matching in the presence of directional shadows cast by scattered objects. Indeed numerous desert-dwelling geckos have been observed to have contrasting bars despite living within an open habitat (Allen et al., 2020).

While not measured in this study, the orientation of the observer’s field of view is also likely to influence the perception of orientations of patterns generated by cast and received shadows from direct lighting (Hancock et al., 2023; Li et al., 2026). Due to limitations in the range of our 3D scanner and the inability to control lighting when viewed from the horizon (the photography tent and shadows from the tripod would have been visible), we opted for the bird’s eye view that is commonly used for photography intended for image analysis of backgrounds and animals for camouflage and signaling experiments. These changes in angle are likely to enhance or reduce the influence of lighting on the background in much the same way as differences in the orientation of light influence countershading on an animal but still averaging to our measured effects (Penacchio et al., 2018).

In conclusion, our study serves both as a showcase for both how 3D scanners can be used alongside image metrics and for how the measured image properties of the environment are directly affected by their geometry and lighting structure with respect to the scale of the target of interest. The increased accessibility of outdoor operable 3D scanners allowed us to effectively measure the local 3D variation within natural scenes and its effects on scene statistics. Here, we present human-color-based modeling of a set of human-scale environments, so we must be cautious about generalization to other species and other habitats. How environmental conditions influence pattern evolution in particular species requires species-specific application of our methods to the color, lighting and geometry of the relevant local surroundings of that species. Future research on animal camouflage strategies should explore the interactions between background geometry and lighting environment on animal pattern contrast, both luminance and chromatic, and orientation, either by comparative pattern analyses of animals that occupy measured habitats or testing predictions from modeling such as ours with visual-search-based experiments with different pattern phenotypes.

## Author Contributions

The initial concept for assessing the interactions between geometry and the orientation of lighting were conceived by JT and carried out by GRAH. GRAH collected color-calibrated photographs and 3D scans. All code was written by JT and GRAH. The manuscript was first written by GRAH, with subsequent edits by all authors. GRAH performed data analyses with guidance from JT and IC.

## Acknowledgements.

Natural Environment Research Council (NERC) GW4+ NE/S007504/1 funded GRAH in a CASE partnership with the Game and Wildlife Conservation Trust (GWCT). JT was funded by a NERC Independent Research Fellowship NE/P018084/1. ICC was funded by Biotechnology & Biological Sciences Research Council grant BB/S00873X/1.

## Data and resource availability

All image and 3D environment measures as well as the R scripts required to perform our analyses are provided within our FigShare data repository (Private Link - > https://figshare.com/s/a0ad09ab601f697fdc7b), along with the ImageJ scripts used to perform them and sample habitat images and 3D scans.

