## supplementary material for "The effects of habitat structure and lighting on object and background appearance"

### Supplementary Material for Hancock et al. "The effects of habitat structure and lighting on object and background appearance"

George R.A. Hancock[1]*, Innes C. Cuthill[2], Jolyon Troscianko[1]

1. Centre for Ecology & Conservation, University of Exeter, Penryn, TR10 9FE, UK

2. School of Biological Sciences, University of Bristol, Tyndall Avenue, Bristol BS8 1TQ, UK

#### Photography Protocol

Given the image dimensions of both the camera photos and the size of the photography tent (1.5m$^3$) the tripod arm had to be modified by attaching an additional telescopic pole to extend the arm >0.75m into the centre of the tent. A counterweight was attached and adjusted to the opposite end of the tripod to prevent the phone camera from dropping. To activate the camera a Bluetooth remote was paired with the ASUS zenfone (ZenFone AR ZS571KL, AsusTek Computer Inc., Taipei, Taiwan) so that it could be used remotely from outside the photography tent and to avoid moving the camera (Figure A1).

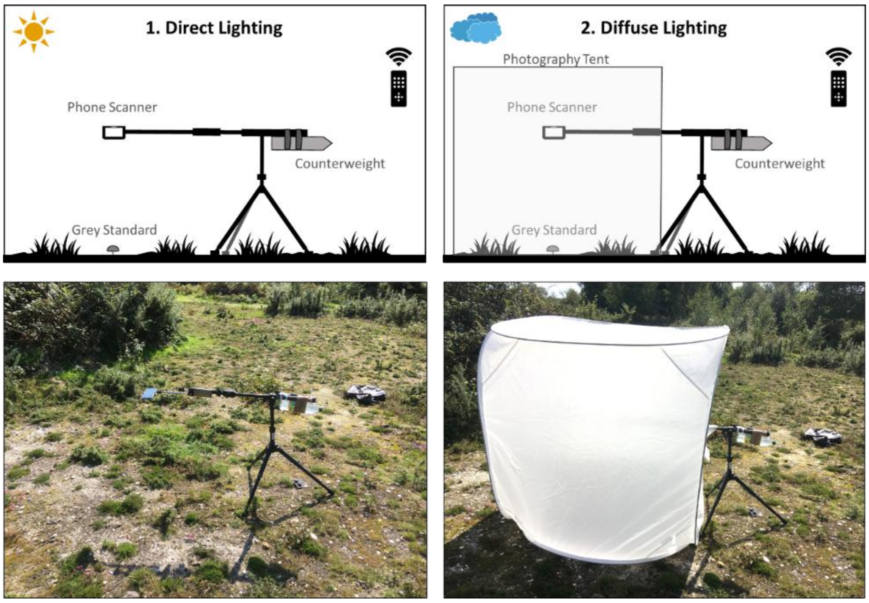


Figure A1. Diagram of habitat photography methods for direct lighting (left) and diffuse lighting (right) conditions.

#### Occlusion Removal

Within vegetated habitats small portions of vegetation overlapped in front of the edges of the gey-standard (Figure A2). To remove them for analyses the sections were highlighted and not used for normalisation measurements. Post-normalisation the occluded vegetation was removed by photo-editing in GIMP 2.10.36

(https://www.gimp.org/), the replaced regions were meticulously sampled from the standard with the clone and paintbrush tool.

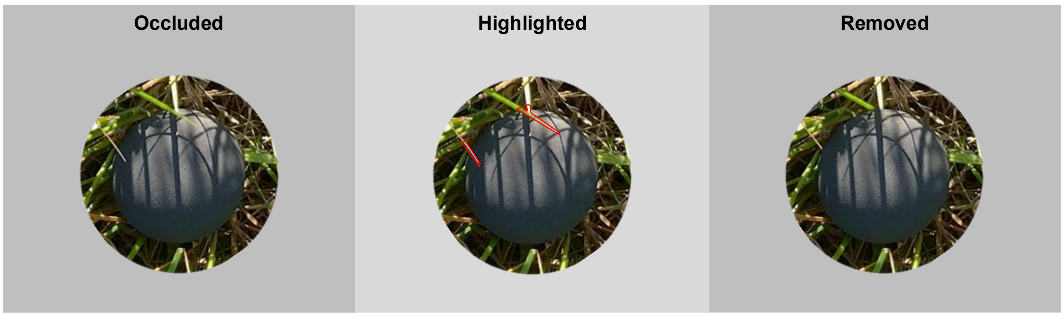


Figure A2. Illustration of steps for removal of occluding vegetation. Occluding vegetation was identified and replaced with samples from the target with the clone tool in GIMP.

### Background Colour Maps

Direct lighting effects the mean colour of the backgrounds not just contrast, direct lighting backgrounds (Mean = are darker and bluer compared to the diffuse backgrounds when considered relative to the grey target (Figure A3).

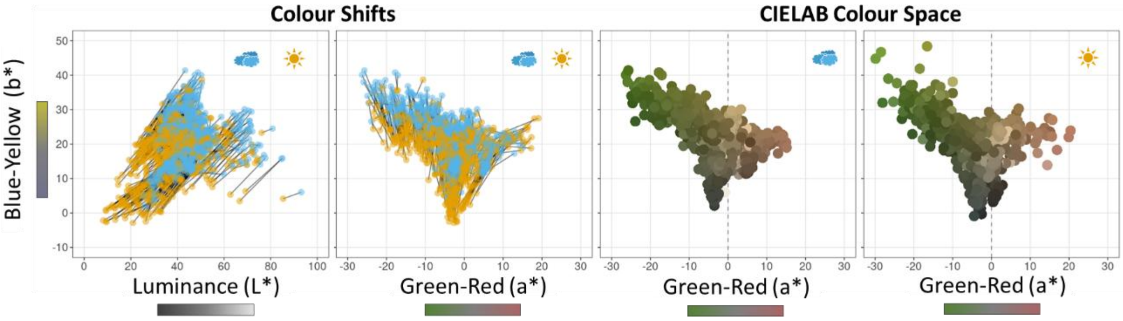


Figure A3. Shifts in colour space between direct and diffuse lighting, left-two panels show comparison between each location in direct and diffuse lighting. The right-two panels show the rendered average CIE LAB colour.

#### Background Measures

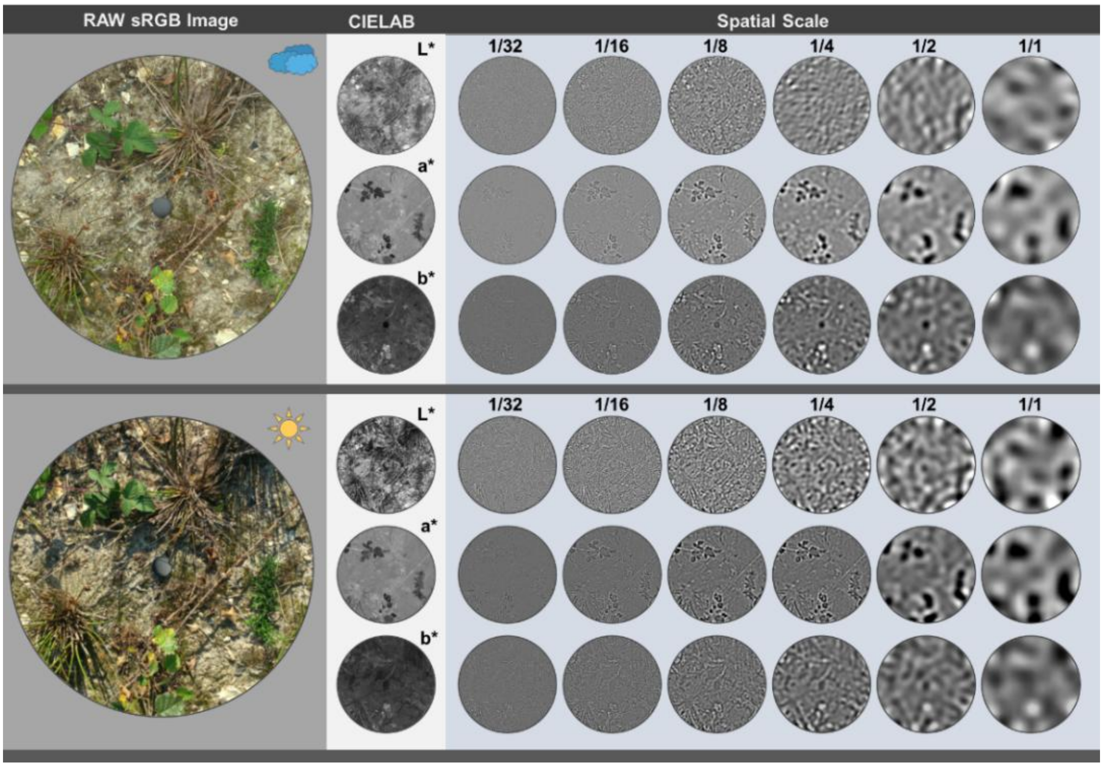


Figure A4. Contrast measures for CIELAB, L* (Luminance), a* (Green-Red) and b* (Blue-Yellow) channels for the same habitat under diffuse (above) and direct (below) lighting. Target is maintained as a scale reference for the figure only, otherwise the target was removed using a mask for analyses.

#### Target Measures

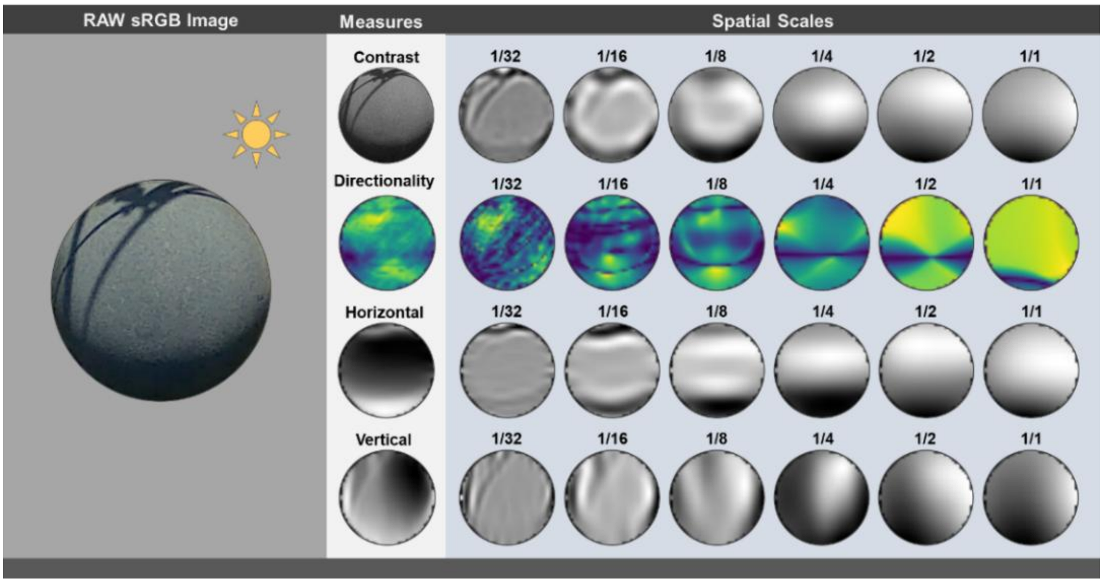


Figure A5. Target object measures for the CIELAB L* (Luminance) channel across the 6 spatial scales. Contrast, directionality, horizontal ratio and vertical ratio. Self-shading appears as a horizontal band while the received shadows from surrounding vegetation create vertical patterns at smaller spatial scales.

### 3D Scans

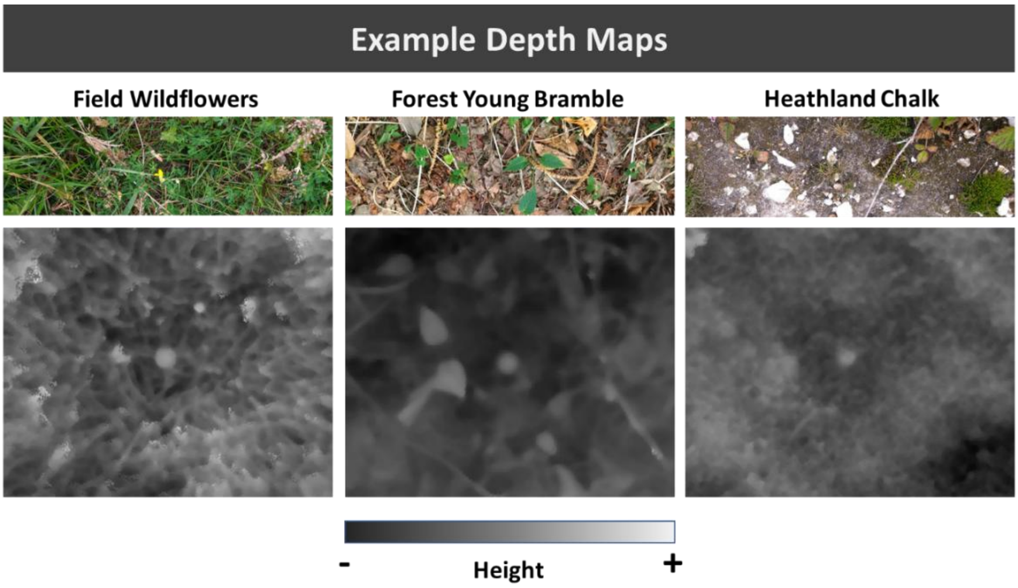


Figure A5. Height map images (below) for three different habitats, field wildflowers, forest young bramble and heathland chalk. More elevated areas are shown as lighter. Against each scene the height and shape of the target make it stick out within the depth map. Habitats are ordered by 3D complexity from high to low, left to right.

### Site Info Table

Table 1. Time of year, location and sun position for the photography session of each habitat.

| Habitat | Year | Month | Day | Time | TimeUTC | County | Lat | Long | SunElevation | SunAzimuth |
|---|---|---|---|---|---|---|---|---|---|---|
| 01_grassWildFlowers | 2020 | 7 | 20 | 18:07:00 | 17:07:00 | Surrey | 51.26512 | -0.26515 | 27.60 | 272.21 |
| 02_grassMowShort | 2020 | 7 | 29 | 09:23:00 | 08:23:00 | Surrey | 51.26523 | -0.26894 | 45.02 | 121.72 |
| 03_woodLeaflitter | 2020 | 7 | 29 | 16:18:00 | 15:18:00 | Surrey | 51.26497 | -0.26917 | 34.17 | 256.61 |
| 04_woodTrail | 2020 | 7 | 30 | 10:36:00 | 09:36:00 | Surrey | 51.266 | -0.26567 | 44.83 | 121.93 |
| 05_grassTrampled | 2020 | 7 | 30 | 14:51:00 | 13:51:00 | Surrey | 51.26575 | -0.26627 | 50.28 | 224.43 |
| 06_grassTopped | 2020 | 7 | 30 | 16:46:00 | 15:46:00 | Surrey | 51.26423 | -0.26787 | 38.47 | 249.57 |
| 07_woodLichen | 2020 | 7 | 31 | 09:38:00 | 08:38:00 | Surrey | 51.26528 | -0.26798 | 36.12 | 107.08 |
| 08_grassMowLong | 2020 | 8 | 3 | 14:07:00 | 13:07:00 | Surrey | 51.26489 | -0.2685 | 54.47 | 202.01 |
| 09_woodBramble | 2020 | 8 | 7 | 09:40:00 | 08:40:00 | Surrey | 51.2672 | -0.26522 | 34.76 | 108.65 |
| 10_woodEdgeGrass | 2020 | 8 | 11 | 11:49:00 | 10:49:00 | Surrey | 51.26457 | -0.26762 | 49.00 | 143.14 |
| 11_heathHeather | 2020 | 9 | 10 | 11:19:00 | 10:19:00 | Surrey | 51.26635 | -0.27593 | 37.38 | 141.89 |
| 12_heathChalk | 2020 | 9 | 14 | 10:09:00 | 09:09:00 | Surrey | 51.2677 | -0.27669 | 29.49 | 127.01 |
| 13_heathPond | 2020 | 9 | 14 | 11:28:00 | 10:28:00 | Surrey | 51.26927 | -0.27905 | 38.62 | 151.90 |
| 14_heathGrass | 2020 | 9 | 15 | 15:30:00 | 14:30:00 | Surrey | 51.26847 | -0.27764 | 31.78 | 226.96 |
| 15_heathFerns | 2020 | 9 | 17 | 15:09:00 | 14:09:00 | Surrey | 51.26655 | -0.27638 | 34.20 | 218.67 |
| 16_heathBog | 2020 | 9 | 17 | 15:47:00 | 14:47:00 | Surrey | 51.26915 | -0.27849 | 31.02 | 226.73 |
| 17_heathErosion | 2020 | 9 | 18 | 15:19:00 | 14:19:00 | Surrey | 51.26654 | -0.27668 | 33.81 | 218.59 |
| 18_heathScrub | 2020 | 9 | 21 | 10:51:00 | 09:51:00 | Surrey | 51.26613 | -0.27726 | 30.86 | 136.78 |
| 19_woodNettle | 2020 | 9 | 22 | 13:13:00 | 12:13:00 | Surrey | 51.26442 | -0.26931 | 38.78 | 182.03 |
| 20_estuaryPebble | 2020 | 10 | 14 | 11:38:00 | 10:38:00 | Cornwall | 50.16881 | -5.09173 | 27.85 | 152.86 |
| 21_estuarySeaweed | 2020 | 11 | 4 | 10:53:00 | 10:53:00 | Cornwall | 50.16887 | -5.09146 | 21.13 | 155.72 |
| 22_estuaryGravel | 2020 | 11 | 4 | 15:24:00 | 15:24:00 | Cornwall | 50.16889 | -5.09156 | 10.27 | 229.99 |
| 23_estuaryVeg | 2020 | 11 | 5 | 10:34:00 | 10:34:00 | Cornwall | 50.16903 | -5.09014 | 20.84 | 155.80 |
| 24_estuarySilt | 2020 | 11 | 5 | 14:33:00 | 14:33:00 | Cornwall | 50.16876 | -5.09108 | 16.57 | 216.64 |
| 25_farmPlough | 2020 | 11 | 26 | 10:10:00 | 10:10:00 | Cornwall | 50.16898 | -5.0888 | 13.44 | 149.50 |
| 26_woodFlower | 2021 | 3 | 19 | 12:32:00 | 12:32:00 | Surrey | 51.26404 | -0.26922 | 38.20 | 186.77 |
| 27_gravelMajority | 2021 | 3 | 29 | 11:52:00 | 11:52:00 | Hampshire | 50.93136 | -1.78465 | 41.82 | 165.93 |
| 28_gravelMix | 2021 | 4 | 2 | 13:10:00 | 12:10:00 | Hampshire | 50.93116 | -1.78485 | 44.13 | 176.31 |

### Background Metrics Table

Table 2. The average L, a, b and z values for each habitat. Shows the mean and std dev for colour (L,a,b) and just std dev for height (z).

| Habitat | Lighting | L* Std Dev | L* Mean | a* Std Dev | a* Mean | b* Std Dev | b* Mean | Z Std Dev |
|---|---|---|---|---|---|---|---|---|
| Field Wildflower | Diffuse | 14.98 | 39.27 | 10.33 | -14.57 | 10.94 | 31.79 | 21.34 |
| Field Wildflower | Direct | 21.45 | 40.65 | 10.4 | -15.56 | 14.73 | 30.75 | 21.34 |
| Field Topped | Diffuse | 12.43 | 44.75 | 9.07 | -16.74 | 10.48 | 35.43 | 14.5 |
| Field Topped | Direct | 15.84 | 39.35 | 8.04 | -15.46 | 11.03 | 31.1 | 14.5 |
| Ferns | Diffuse | 14.98 | 44.48 | 10.12 | -17.43 | 11.8 | 34.7 | 45.49 |
| Ferns | Direct | 19.26 | 40.72 | 9.65 | -16.01 | 12.8 | 28.84 | 45.49 |
| Forest Nettle | Diffuse | 16 | 49.1 | 13.68 | -17.5 | 10.71 | 32.34 | 41.94 |
| Forest Nettle | Direct | 20.2 | 45.48 | 13.46 | -17.19 | 11.76 | 28.36 | 41.94 |
| Forest Flowers | Diffuse | 17.32 | 44.4 | 13.76 | -14.51 | 12.45 | 33.75 | 27.24 |
| Forest Flowers | Direct | 21.48 | 36.63 | 12 | -14.05 | 13.73 | 27.41 | 27.24 |
| Bog Grass | Diffuse | 15.12 | 43.24 | 9.74 | -16.15 | 10.46 | 31.97 | 21.13 |
| Bog Grass | Direct | 19.8 | 37.35 | 9 | -15.36 | 11.82 | 26.13 | 21.13 |
| Scrub Grass | Diffuse | 11.8 | 45.95 | 9.4 | -13.78 | 7.85 | 28.83 | 6.05 |
| Scrub Grass | Direct | 14.97 | 41.04 | 8.28 | -13.34 | 8.43 | 24.63 | 6.05 |
| Field Eroded | Diffuse | 10.77 | 50.48 | 8.28 | -9.66 | 7.53 | 29.69 | 8.78 |
| Field Eroded | Direct | 16.05 | 45.14 | 8.1 | -8.55 | 7.78 | 25.33 | 8.78 |
| Heath Heather | Diffuse | 13.59 | 38.95 | 9.94 | -9.09 | 10.4 | 25.61 | 17.73 |
| Heath Heather | Direct | 17.93 | 28.99 | 9.65 | -7.86 | 10.42 | 17.57 | 17.73 |
| Pond Bank | Diffuse | 13.28 | 42.47 | 7.62 | -5.67 | 8.75 | 19.18 | 11.74 |
| Pond Bank | Direct | 18.8 | 35.11 | 6.9 | -6.1 | 9.67 | 13.7 | 11.74 |
| Lawn Uncut | Diffuse | 12.21 | 47.09 | 6.89 | -5.05 | 6.87 | 26.76 | 7.42 |
| Lawn Uncut | Direct | 15.53 | 42.24 | 6.94 | -5.23 | 7.18 | 21.77 | 7.42 |
| Lawn Cut | Diffuse | 11.32 | 45.96 | 8.81 | -11.05 | 8.86 | 32.56 | 5.84 |
| Lawn Cut | Direct | 16.01 | 36.13 | 8.68 | -9.07 | 10.16 | 25.48 | 5.84 |
| Estuary Mud Flat | Diffuse | 10.78 | 46.75 | 3.75 | -2.85 | 4.9 | 14.63 | 5.37 |
| Estuary Mud Flat | Direct | 17.93 | 36.37 | 3.46 | -3.66 | 9.06 | 6.14 | 5.37 |
| Estuary Gravel | Diffuse | 18.83 | 47.42 | 4.6 | 0.11 | 7.86 | 17.8 | 9.69 |
| Estuary Gravel | Direct | 23.59 | 37.69 | 3.92 | -0.98 | 10.04 | 9.96 | 9.69 |
| Estuary Plants | Diffuse | 19.99 | 41.71 | 8.44 | -5.89 | 11.68 | 26.72 | 13.71 |
| Estuary Plants | Direct | 22.83 | 34.52 | 7.4 | -6.08 | 12.32 | 20.95 | 13.71 |
| Gravel Succession | Diffuse | 15.67 | 47.05 | 8.93 | -5.27 | 9.87 | 27.01 | 3.86 |
| Gravel Succession | Direct | 18.19 | 44.14 | 8.82 | -4.08 | 9.51 | 23.68 | 3.86 |
| Heath Eroded | Diffuse | 14.5 | 68.84 | 4.62 | -0.41 | 5.47 | 19.22 | 11.4 |
| Heath Eroded | Direct | 19.06 | 66.39 | 4.47 | -0.34 | 6.14 | 18.6 | 11.4 |
| Estuary Seaweed | Diffuse | 16.63 | 32.18 | 2.3 | -3.53 | 4.84 | 8.17 | 13.5 |
| Estuary Seaweed | Direct | 21.27 | 21.88 | 2.36 | -2.74 | 7.17 | 4.18 | 13.5 |
| Estuary Pebbles | Diffuse | 14.81 | 41.76 | 3.81 | -4.83 | 6.43 | 14.2 | 15.02 |
| Estuary Pebbles | Direct | 22.98 | 32.15 | 3.57 | -4.49 | 9.01 | 8.04 | 15.02 |
| Gravel | Diffuse | 23.31 | 70.24 | 6.16 | 0.78 | 11.19 | 26.31 | 5.49 |
| Gravel | Direct | 28.75 | 60.11 | 5.83 | 1.24 | 12.17 | 19.17 | 5.49 |
| Forest Succession | Diffuse | 11.75 | 47.23 | 7.03 | 0.9 | 5.86 | 24.61 | 7.59 |
| Forest Succession | Direct | 18.81 | 40.2 | 8.44 | 0.69 | 8.81 | 22.11 | 7.59 |
| Heath Chalk | Diffuse | 16.65 | 45.55 | 6.7 | -4.38 | 9.12 | 19.08 | 8.44 |
| Heath Chalk | Direct | 19.78 | 41.7 | 6.19 | -3.8 | 8.71 | 16.28 | 8.44 |
| Heath Succession | Diffuse | 16.46 | 52.77 | 7.69 | -3.44 | 8.49 | 22.84 | 15.21 |
| Heath Succession | Direct | 20.01 | 46.33 | 7.2 | -3.87 | 10.96 | 16.7 | 15.21 |
| Forest Bramble | Diffuse | 17.31 | 49.63 | 9.33 | 2.11 | 6.44 | 20.49 | 14.64 |
| Forest Bramble | Direct | 22.8 | 47.94 | 11.57 | 3.92 | 8.57 | 20.09 | 14.64 |
| Field Ploughed | Diffuse | 13.9 | 37.76 | 2.61 | 5.35 | 4.58 | 13.78 | 20.74 |
| Field Ploughed | Direct | 21.96 | 21.39 | 3.34 | 1.07 | 7.85 | 5.27 | 20.74 |
| Forest Trail | Diffuse | 14.28 | 51.49 | 5.31 | 9.11 | 4.84 | 20.9 | 9.57 |
| Forest Trail | Direct | 21.87 | 49.99 | 8.61 | 8.01 | 7.02 | 18.18 | 9.57 |
| Forest Leaflitter | Diffuse | 16.57 | 49.01 | 6.46 | 10.05 | 6.17 | 22.06 | 11.27 |
| Forest Leaflitter | Direct | 23.53 | 39.74 | 7.38 | 6.18 | 8.25 | 16.7 | 11.27 |
| Forest Edge | Diffuse | 16.28 | 52.72 | 7.1 | 2.93 | 6.01 | 22.88 | 7.06 |
| Forest Edge | Direct | 21 | 50.1 | 7.85 | 3.2 | 7.43 | 21.76 | 7.06 |
